# Novel Calcin peptides from Scorpion Venom Exhibit Humidity-Dependent Expression and Ryanodine Receptor Activation

**DOI:** 10.1101/2025.09.17.676761

**Authors:** Xiaoyu Hua, Zhixiao Yang, Li Xiao, Songyu Gao, Fengling Yang, Bilin Tao, Yi Wang, Carmen R. Valdivia, Wei Chen, Marina Pozzolini, Lingxin Chen, Héctor H Valdiva, Liang Xiao

## Abstract

Scorpions are venomous animals of considerable pharmacological potential, traditionally used in East Asia to treat cardiovascular and autoimmune diseases. However, geographical variations in the expression of their venom components remain poorly understood. We performed transcriptome profiling on scorpion samples collected from eight distinct climatic and environmental regions in East Asia, we successfully identified two distinct scorpion species: *Uroctonus mordax* and *Mesobuthus martensii*. Furthermore, we identified two novel calcins from the venoms of these species, with their expression levels showing a significant correlation to environmental humidity. Functional assays revealed that these calcins differentially potentiate [³H]ryanodine binding to RyR isoforms and induce a persistent subconductance state, suggesting distinct mechanisms of action. Our findings suggest biogeographical variation in venom composition between species and suggest that environmental humidity as a key factor influencing calcin expression. These results expand the structural diversity of the calcin family and provide a mechanistic basis for the medicinal application and ecological adaptation of scorpion venom.

## 1. Introduction

Scorpions are among the most medically significant venomous animals, with their venom comprising a complex cocktail of ion channel-modulating toxins and inflammatory components(Ma, Zhao et al., 2009, Ruiming, Yibao et al., 2010). Envenomation represents a critical public health burden, affecting over two billion people globally and causing more than one million stings annually(Pucca, Fry et al., 2021, Sabatier, 2011). The most severe clinical manifestations include acute heart failure, cardiogenic shock, and pulmonary edema, with a mortality rate reaching 0.27%(Abroug, Souheil et al., 2015, Fereidooni, Shirzadi et al., 2023). The escalating impact of scorpion stings underscores the urgent need for more effective therapeutic and preventive strategies(Bergman, 1997).

Despite their toxicity, scorpions have been used for millennia in traditional Chinese medicine and throughout East Asia to treat cardiovascular and autoimmune diseases(Mohammadi Bavani, Rafinejad et al., 2021, Zeng, Luo et al., 2006). This paradox stems from the venom’s dual nature: it contains not only harmful toxins but also short-chain peptides (typically 20-80 amino acids stabilized by three or four disulfide bridges) that exhibit high specificity for ion channels(Abroug, Ouanes-Besbes et al., 2019, Ding, Chua et al., 2014, Uzair, Bint et al., 2018). By modulating electrophysiological activity through channel blockade or alterations in gating kinetics, these peptides induce pharmacologically beneficial effects including analgesia, anti-inflammation, and detoxification(Bouhaouala-Zahar, Ben Abderrazek et al., 2011, Lin King, Emrick et al., 2019). These mechanisms render them promising candidates for treating neurological and inflammatory disorders such as epilepsy, rheumatoid arthritis, and chronic pain(Bhavya, Francois et al., 2016, Gao, Liang et al., 2021). Thus, the same venom responsible for scorpions’ lethality also represents a valuable resource for drug discovery. Among these, peptides targeting ryanodine receptors (RyRs), key calcium release channels in sarcoplasmic reticulum, have emerged as particularly important for their cardiotoxic and potential cardioprotective effects(Smith, Vetter et al., 2013, Vargas-Jaimes, Xiao et al., 2017, Zhu, Zamudio et al., 2004).

The therapeutic potential and pathological severity of scorpion venom are intimately linked to its compositional diversity, which varies not only among species but also within species across different geographic and climatic regions(Chowell, Hyman et al., 2005, Rodríguez de la Vega, Schwartz et al., 2010, Santibáñez-López, Graham et al., 2019). Environmental factors such as temperature, humidity, and altitude are increasingly recognized as key drivers of venom variability, potentially enhancing ecological adaptability and venom efficacy(Adams, Marais et al., 2016, Agourram, Zegrari et al., 2024, Elmourid, Boussaa et al., 2023). For instance, calcins-a family of peptides targeting RyRs-exhibit sequence variation, differential expression, and functional divergence across species and populations, suggesting local environmental adaptation(Li, Wang et al., 2023, Xiao, Gurrola et al., 2016). However, direct evidence linking specific ecological factors to scorpion venom variation remains limited, particularly at the transcriptomic level and across wide climatic gradients.

To address this gap, we conducted a broad-scale transcriptomic analysis of venom glands from scorpions collected across eight distinct climatic regions in East Asia. This study was designed to identify novel calcin peptides, quantify their region-specific expression levels, and examine potential correlations between toxin expression and key environmental variables. Our findings offer novel insights into the ecological mechanisms shaping venom diversity and facilitate the discovery of environmentally adapted bioactive peptides with therapeutic promise.

## 2. Materials and Methods

### 2.1 Collection of scorpion and toxin samples

Scorpion samples (mixed male and female) were collected from 8 regions in East Asia, including AH (Anhui, Hefei), SX (Shanxi, Taiyuan), XJ (Xinjiang, Hetian), QH (Qinghai, Xining), Tibet (Linzhi), LN (Liaoning, Fushun), NX (Ningxia, Wuzhong) province of China, and Myanmar. Due to the preciousness and collection difficulty of wild scorpion samples, the test samples in this study were divided into two batches. The first batch was scorpions from AH, LN, MM, and Tibet regions, with 30 scorpions collected from each region and used to construct transcriptomic libraries (n=5). The second batch was scorpions from SX, NX, QH, and XJ regions, with only 10 scorpion samples collected from each region. The pooled sequencing method was used to construct libraries for subsequent determination. Additionally, 1 mL of scorpion venom were harvested using electrical stimulation and stored at -80°C prior to analysis.

### 2.2 Collection of climate and environmental data

The climate and environmental data collected spanned from 2013 to 2023, encompassing measurements of annual average temperature, sunshine duration, humidity, and rainfall. The data of seven regions in China were obtained from the National Climatic Data Center (https://data.cma.cn/). The climate data for the Myanmar region is sourced from the WorldClim database (https://www.worldclim.org/data/worldclim21.html).

### 2.3 Total RNA extraction and sequencing

Total RNA was extracted by the Trizol (Invitrogen, CA, USA) method(Rio, Ares et al., 2010). The total RNA quantity and purity were analysis of Bioanalyzer 2100 and RNA 1000 Nano LabChip Kit (Agilent, CA, USA) with RIN number >7.0. Poly(A) RNA is purified from total RNA (5 µg) using poly-T oligo-attached magnetic beads using two rounds of purification. Following purification, the mRNA is fragmented into small pieces using divalent cations under elevated temperature. Then the cleaved RNA fragments were reverse-transcribed to create the final cDNA library in accordance with the protocol for the mRNA-Seq sample preparation kit (Illumina, San Diego, USA), the average insert size for the paired-endlibraries was 300 bp (±50 bp). And then we performed the paired-end sequencing on an Illumina Novaseq™ 6000 at the (LC Sceiences, USA) following the vendor’s recommended protocol.

Cutadapt and perl scripts in house were used to remove the reads that contained adaptor contamination, low quality bases and undetermined bases. Then sequence quality was verified using FastQC, including the Q20, Q30 and GC-content of the clean data. All downstream analyses were based on clean reads of high quality. The transcriptome expression quantification software RSEM was used to align the clean reads of each sample to the reference sequence using the spliced transcript sequences as a reference(Li & Dewey, 2011). Then, the number of reads aligned to each gene in each sample was counted, the FPKM values at the gene level and transcript level were calculated, and the expression differences at the gene level and transcript level were calculated to screen the differentially expressed genes between samples or groups. Trinity 2.4.0 was used to assemble high-quality sequences following quality control(Grabherr, Haas et al., 2011). A strategy of co-assembling all samples was adopted to normalize the samples, ultimately generating unigenes.

### 2.4 Bioinformatics analysis

All unigenes were aligned with the NR database (http://www.ncbi.nlm.nih.gov/), GO (http://www.geneontology.org), SwissProt Database (http://www.expasy.ch/sprot/) and KEGG database (http://www.genome.jp/kegg/)(Conesa, Götz et al., 2005, Okuda, Yamada et al., 2008). R package edgeR selection(Robinson, McCarthy et al., 2010) with several comparative parameters including log2Ratio > 1 or log2Ratio < -1, p values < 0.01, and q < 0.05. GO and KEGG analyses of differentially expressed functional genes were performed by internal Perl script. GO analysis was performed with three terms, namely, biological process (BP), cellular component (CC) and molecular function (MF). The signaling pathways were analyzed by comparison with the KEGG database. The ORFfinder online service (https://www.ncbi.nlm.nih.gov/orffinder) was used to translate the nucleotide sequences into amino acid sequences. Multiple sequence alignment was performed using BLAST and BioEdit software (version: 7.2.5)(Hall, 1999). Blast was used for manual editing of the obtained sequences, adjustment of alignment parameters, and other detailed processing. The phylogenetic tree was constructed by the adjacency method with 1000 bootstrap replication sets using MEGA7(Kumar, Stecher et al., 2016).

### 2.5 Toxin screening

The local BLAST comparison was conducted between the scorpion venom gland transcriptome and the Uniprot database (https://www.uniprot.org/) using a threshold of E-value less than 10^−5^ to identify toxin-related genes at the transcriptome level. Each identified toxin was subsequently classified based on its expression level.

### 2.6 Cytotoxicity assay

H9C2 cells were grown in DMEM medium containing 10% fetal bovine serum and 1% antibiotics, cultured at 37℃, 5% CO_2_.The viability was assessed using the CCK8 method(Robinson et al., 2010). The cells were cultured in a 96-well plate at a density of 5000 cells per well. Following a 24 h incubation period, the cells were exposed to various concentrations of scorpion venom (0-13 mg/mL, n = 4) for a duration of 2 h. Then, the supernatant was discarded, and 90 µL of DMEM (BasalMedia, China) along with 10 µL of CCK-8 reagent (Target, USA) were added to each well. The plates were then incubated at 37℃ for 2 h. The change in absorbance was measured at 450 nm using a microplate reader (BioTek, USA). The formula for calculating cell survival rate was as follows: Viability (%) = (OD _scorpion venom_ - OD _Background_)/ (OD _Control_ - OD _Background_) × 100%. In this context, OD _scorpion venom_ denotes the optical density observed in the test sample, OD _Background_ represents the optical density recorded when using only DMEM without cell, and OD _Control_ signifies the optical density associated with the negative control which consists of DMEM.

### 2.7 Metalloprotease activity determination

The metalloproteinase activity of scorpion venom was assessed using the azocasein method^(Robinson^ ^et^ ^al., 2010)^. Using PBS as the negative control and batimastat inhibitor (1 µM) as the positive control. Add the scorpion venom from different regions at a concentration of 10 mg/mL to the buffer solution containing Tris-HCl, PBS, NaCl, CaCl_2_, and azocasein (Sigma, USA) (pH 8.8). The mixture was incubated in a water bath at 37°C for 90 min. After incubation, 200 μL of 0.5 M trichloroacetic acid (TCA) (Ronoen, China) was added to each sample and allowed to stand at room temperature for 30 min. Subsequently, the samples were centrifuged at 10,000×*g* for 10 min. PBS served as the negative control group and batimastat as the positive control group. From each sample, 100 μL of the supernatant was collected and mixed with 100 μL of 0.5 M NaOH (Amresco, USA). The absorbance of the resulting mixtures was measured at 450 nm, with each experiment performed four times.

### 2.8 PLA_2_ activity determination

4-Nitro-3-octanoxybenzoic acid (NOBA, Sigma, USA) was prepared to a concentration of 1 mg/mL in acetonitrile and mixed with Tris-HCl (pH 8.8) buffer to achieve a final buffer concentration of 50 mM, supplemented with 10 mM CaCl₂ and 100 mM NaCl. To a 96-well culture plate, 25 µL of scorpion venom (with concentrations ranging from 0 to 10 mg/mL, n=6) was added, followed by the addition of 200 µL Tris-HCl and 25 µL NOBA. The plates were then incubated for 1 h at 37°C. PBS served as the negative control group. Subsequently, absorbance was measured at OD_405_ nm using a microplate reader (BioTek, USA). The PLA_2_ activity was determined by subtracting the optical density of the negative control group from that of the scorpion toxin.

### 2.9 Chemical synthesis and identification

All calcin peptides were synthesized through QYAOBIO (ChinaPeptides Co., Ltd.). Calcin linear peptides were first synthesized by the Fmoc–amino acids solid phase synthesis method, then cut with 95% TFA solution for 3 h at normal temperature, extracted with 5% acetic acid, and dried under vacuum. A folding buffer solution (20 mM Na_2_HPO_4_, 0.1 M NaCl, 5 mM GSH and 0.5 mM GSSG) was added to cyclize the molecules into disulfide bonds under natural oxidation conditions. The pH was adjusted to 7.9 with 1 M NaOH and oxidized at normal temperature for 4 h.

After oxidation of the disulfide bond, the pH of the polypeptide solution was adjusted to 3.0 with formic acid, followed by purification (purity ≥95%) using a C_18_ column (100-5-C_18_, 4.6 mm×250 mm, 5 µm, Kromasil). The purification was performed using an Agilent LC1200 high-performance liquid chromatograph with a linear gradient elution from 0 to 60% over 60 min. The mobile phases were Buffer A (water containing 0.1% trifluoroacetic acid) and Buffer B (acetonitrile containing 0.1% trifluoroacetic acid). The flow rate was set at 1 mL/min, and the detection wavelength was 220 nm.

Mass spectrometry identification was performed in electrospray ionization (ESI) positive ion mode, with data acquisition ranging from m/z 100 to 2000, a scanning time of 0.5 seconds, and an interval time of 0.1 seconds. Data collection and processing were conducted using MassLynx software. The MaxEnt1 algorithm was applied for deconvolution of multiply charged ions to obtain the molecular mass information of peptides, which was confirmed by comparison with the theoretical calcin molecular weight. All mass spectrometry data were acquired on a Waters zq2000 mass spectrometer.

### 2.10 Heavy Sarcoplasmic Reticulum preparation

Heavy SR (Heavy Sarcoplasmic Reticulum, HSR) was extracted and purified from rabbit skeletal muscle following a previously described method with small modifications(Meissner, 1984). Approximately 300 g of frozen rabbit skeletal muscle was blended for 120 s in ∼500 mL of 0.3 M sucrose and 20 mM HEPES pH 7.2. The minced tissue was placed in Potter-Elvehjem homogenizer, and the above buffer containing protease inhibitor cocktail was added for 60 s at 500 r/min. The homogenate was centrifuged at 4°C for 20 min at 4,000×*g*. The obtained supernatant was filtered through two layers of cheesecloth and spun at 4 °C for 20 min at 8,000×*g*. The new supernatant collected again was slowly added to solid KCl, and a final concentration of 0.6 M was obtained. After the mixture was stirred for 30 min at 4 °C, the supernatant was centrifuged at 100,000×*g* for 30 min. Then, the pellet was collected and resuspended in 50 mL of 0.3 M sucrose, 0.4 M KCl, 5 mM NaPIPES, pH 6.8 (10% sucrose buffer), and placed on a discontinuous sucrose gradient consisting of 11 mL of 20% (w/v), 11 mL of 30%, and 7 ml of 40% sucrose. After centrifugation for 16∼17 h at 120,000×*g*, membranes at sucrose interfaces between 30% and 40% were collected and diluted with 0.4 M KCl, 5 mM NaPIPES, pH 6.8, followed by centrifugation at 100,000×*g* for 30 min. The pellets were resuspended in 0.3 M sucrose, 0.1 M KCl, and 5 mM NaPIPES, pH 6.8, quickly frozen and stored at -80°C before use. The protein concentration was determined by the Bradford method.

### 2.11 [^3^H]ryanodine binding assay

[^3^H]ryanodine binding was carried out following a procedure that was previously described(Schwartz, Capes et al., 2009, Xu & Narayanan, 2000). Binding mixtures were prepared containing 20 µg of protein from rabbit skeletal HSR. To achieve concentration-dependent binding, calcins were directly diluted in incubation buffer (0.2 M KCl, 10 µM CaCl_2_, 10 mM HEPES, pH 7.4). Peptide concentrations were quantified using the UV module of a Nanodrop spectrophotometer (Concentration (mg/mL) =(A_235_-A_280_)/2.51). We corrected the concentration using Nanodrop (Abs at 220 nm, ∼3 kDa peptide as the standard). The final concentration ranges for calcins were 0.987∼987 nM and 0.387∼387 nM, respectively. The Ca^2+^ sensitivity curves were generated in a buffer containing 0.2 M KCl, 20 mM HEPES (pH 7.4), 5 nM [^3^H]ryanodine (PerkinElmer, USA), 1 mM EGTA, and the appropriate amount of CaCl_2_ to achieve a free [Ca^2+^] range of 10 nM to 1,000 µM. The Ca^2+^/EGTA ratio for these solutions was calculated using MaxChelator (WEBMAXCLITE v1.15; pH = 7.4, Ionic = 0.2 N, Temperature = 37°C). Critically, the actual free [Ca²⁺] in representative buffers across the range was experimentally verified using a calcium-specific ion-selective electrode probe (Themo Scientific). Duplicates of samples (100 µL for each) were prepared and used for binding reactions, which were incubated for 2 h at 37 °C. The resulting reaction mixtures were filtered through Whatman GF/B filters (Whatman, Clifton, NJ, USA) and washed twice with 5 mL of Milli-Q water in a Brandel M24-R Harvester. Nonspecific binding was determined in the presence of 20 μM unlabeled ryanodine (2153770, MP Biomedicals). [^3^H]ryanodine binding was measured by liquid scintillation using BioSafe II counting cocktail (RPI Research) in a Beckman LS6500 counter.

### 2.12 Single-channel recording

Single-channel recordings of rabbit skeletal RyR1 integrated into planar lipid bilayers were carried out according to a previously established method(Uehara, Murayama et al., 2017, Xiao et al., 2016). In brief, the experimental setup consisted of two chambers, namely, the cis and trans chambers. The cis chamber represented the cytosolic side and was maintained at virtual ground with the reference electrode. The trans chamber represented the luminal side and contained the voltage command electrode. Single channels were recorded in a nonsymmetrical system, where the cis chamber contained 500 mM CsHSO_4_ and 20 mM MOPs, while the trans chamber contained 50 mM CsHSO_4_ and 20 mM MOPs. HSR vesicles were introduced into the cis chamber, following which channel recordings were obtained at the holding potentials (-20 mV) both before and after the addition of calcins and mutants. Electrical signals were low-pass filtered at 1∼5 kHz and digitized at a rate of 4 kHz by using a Digi-data 1440A AD/DA interface.

### 2.13 Structural analysis

Nine calcin sequences named Imperacaclin (P59868), Opicalcin1 (P60252), Opicalcin2 (P60253), Maurocalcin (P60254), Hemicalcin (A0A1L4BJ42), Vejocalcin (P0DPT1), Urocalcin (L0GBR1), Intrepicalcin (P0DM30) and Hadrucalcin (B8QG00) were identified using the two novel calcins with BLAST search (http://blast.ncbi.nlm.nih.gov/Blast.cgi). Multiple sequence alignment tool Clustal Omega was used to align the calcin sequences and analyze their identity.

RyR1 model were downloaded from the PDB database (https://www.rcsb.org, PDB ID: 7T65(Yuchi, Yuen et al., 2015)). Functional determination of calcin’s hydrophilicity was performed using an online peptide calculator (http://www.chinapeptides.com/tool.php?isCalu=1). AlphaFold2 was used for the prediction of spatial structures of calcins and mutants (https://alphafold.ebi.ac.uk/)(Jumper, Evans et al., 2021). After obtaining the receptor and ligand structure models, GRAMM-X (http://gramm.compbio.ku.edu/) was used for the docking of RyRs with calcins, and interaction residues were calculated using PDBePISA (https://www.ebi.ac.uk/msd-srv/prot_int/)(Paxman & Heras, 2017, Tovchigrechko & Vakser, 2006). The molecular models were visualized and the interactions between ligands and receptors were comprehensively analyzed using PyMOL(Delano & Bioinformatics, 2002) (version 2.5.2). The molecular dynamics were simulated by Gromacs (version 2019.6) under the condition of periodization at room temperature and pressure, and the force field applied for further analysis was Amber. This force field was used to determine the intermolecular interactions during the MD simulation process. The output parameters (RMSD, SASA, Rg, etc.) were used for subsequent analysis(Van Der Spoel, Lindahl et al., 2005).

### 2.14 Statistical Analysis

Statistical analysis was performed using GraphPad Prism software (version 10.1.2). All data are presented as the mean ± standard error of the mean (SEM). Prior to parametric testing, the normality of distribution was confirmed using the Shapiro-Wilk test, and homogeneity of variances was verified with Levene’s test. For multiple comparisons among groups (e.g., venom treatments from regions AH, LN, SX, NX, and control), one-way ANOVA was applied followed by Tukey’s honestly significant difference (HSD) post hoc test for pairwise comparisons. Differences were considered statistically significant at p<0.05. Significant differences are indicated using lowercase letters (a, b, c, etc.). Groups that share a common letter are not significantly different from each other, whereas groups that do not share a letter differ significantly (p<0.05).

Correlation and linear regression analyses were conducted in the R statistical environment (version 4.5.1). The relationship between environmental humidity and unigene number and toxin expression level was evaluated. The Pearson’s product-moment correlation coefficient was computed along with its 95% confidence interval and associated p-value to quantify the strength and statistical significance of the linear association. A simple linear regression model was then fitted to the data, and the goodness-of-fit was indicated by the coefficient of determination. The regression line along with the 95% confidence interval band was visualized on scatter plots using the ggplot2 package (version 3.3.6). [^3^H]ryanodine binding data were fitted using nonlinear regression analysis in Origin 2021b software (Origin Lab). Hill’s equation was used to determine maximum specific binding and Kd. Data are presented as means ± SD, with number of experiments indicated in figure legends. Statistical significance was determined at p < 0.05.

## 3. Results

### 3.1. Transcriptome assembly, species identification and correlation with climatic factors

To investigate the geographical variation of scorpion venom, we collected samples from multiple regions across China (including AH, LN, Tibet, SX, NX, XJ, and QH) and Myanmar (MM), the latter representing a tropical monsoon climate. These regions encompass a wide geographical gradient and are characterized by diverse climatic conditions, particularly in annual temperatures (Fig 1A). Transcriptomic profiling of scorpions from eight distinct regions revealed a clear phylogenetic divergence into two species: *Uroctonus mordax* (*U. mordax*) and *Mesobuthus martensii* (*M. martensii*) (Fig 1B). Our sequencing yielded a total of 243,528 transcripts and 146,510 unigenes. The high-quality of the data was evidenced by a high proportion of valid data (89.71 ∼ 95.58%) and stable GC content (34.82 ∼ 44.84%) across all samples. Among the assembled unigenes, 42,079 (28.72%) and 39,315 (26.83%) were successfully annotated against the GO and KEGG databases, respectively (Fig. 1C and D, Fig. S1). As scorpions are ectothermic, we hypothesized that environmental factors such as temperature sunlight exposure, and humidity would significantly influence their physiology and gene expression^(Lourenço, Ythier et al., 2008)^. We compiled climatic data from each region over a decade (2013-2023) and correlated them with our transcriptomic findings. Based on average annual rainfall and humidity, we classified AH, MM, LN and Tibet as high-humidity regions, and SX, QH, NX and XJ as low-humidity regions (Fig S2). A positive correlation was observed between regional humidity levels and the number of unigenes assembled in eight regions (Fig 1E and F).

**Fig. 1.**
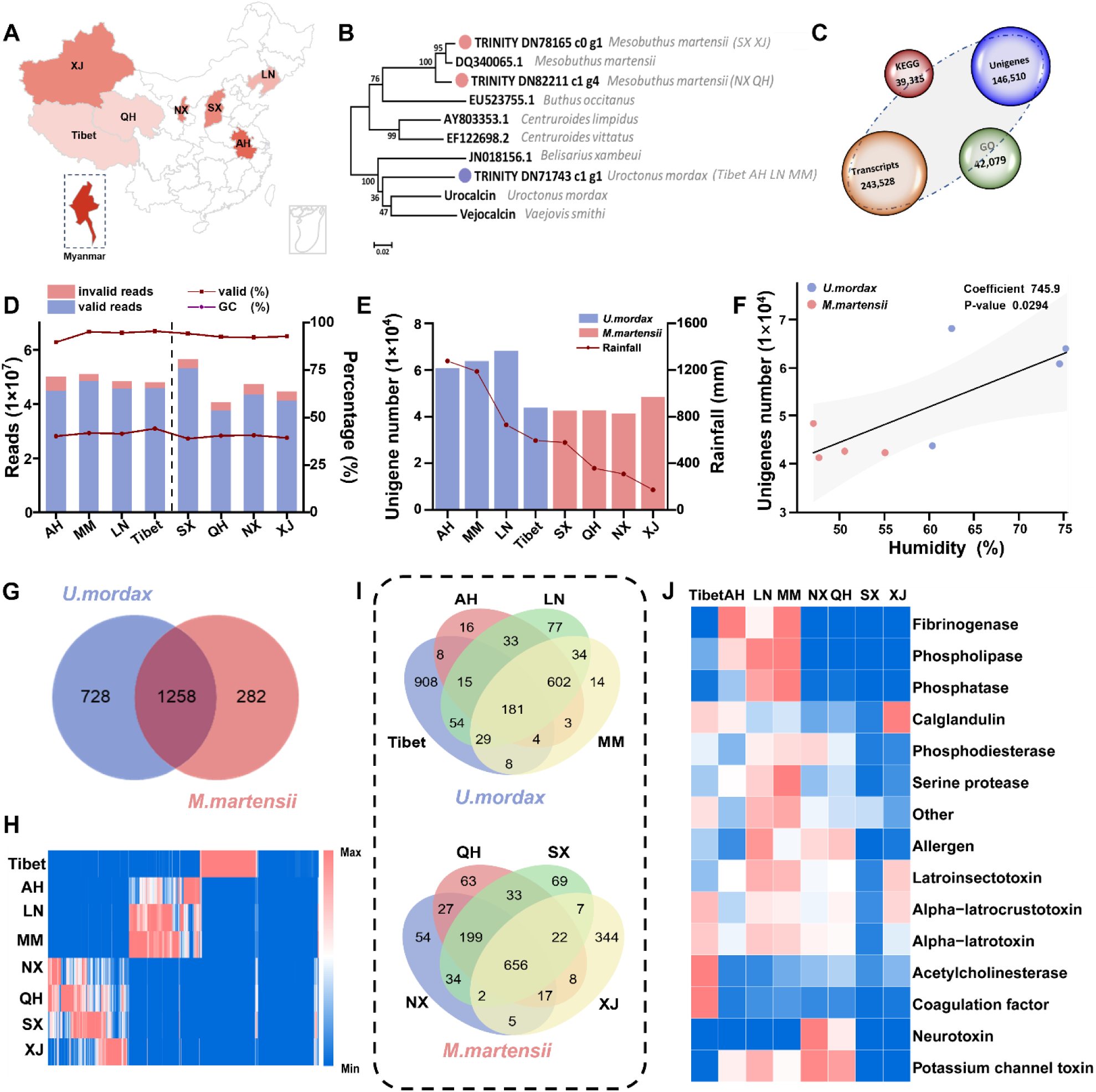
Comparison of environmental factors in 8 East Asian regions and transcriptome identification of scorpions. **(A)** The distribution regions of 8 East Asian scorpions, including Anhui (AH), Shanxi (SX), Xinjiang (XJ), Qinghai (QH), Tibet Liaoning (LN), Ningxia (NX) Province of China, and Myanmar (MM); **(B)** Phylogenetic analysis with the COI sequences of scorpions (red represents *M. martensii* and blue represents *U. mordax*); **(C)** Number of transcripts, unigenes and the number of unigenes annotated in GO and KEGG databases; **(D)** The distribution of reads and GC content among all samples; **(E)** The comparison of the number of unigenes and rainfall in scorpion samples from 8 regions; **(F)** Scatter plot showing the regression line between humidity and the number of unigenes in 8 regions (p<0.05, confidence interval is 95%); **(G)** The number of screened toxins in *U. mordax* and *M. martensii*; **(H)** Heat map of the expression level of toxins in 8 regions; **(I)** The number of screened toxins in the 8 regions of *U. mordax* and *M. martensii*; **(J)** Heat map of the expression level of crucial toxins in 8 regions.

Screening against the Uniprot toxin database identified 2,268 toxin-related sequences. While 1,258 toxins were shared between the two species, *U. mordax* possessed a greater number of unique toxins (728) compared to *M. martensii* (282) (Fig. 1G). Although the expression trends of toxins annotated in the same species were generally consistent across different regions (Fig. 1H), some regions, such as Tibet and XJ still exhibited the specific expression of certain toxins. The four *U. mordax* populations shared 181 common toxins, whereas the four *M. martensii* populations shared 656. Notably, Tibet and XJ exhibited the highest numbers of region-specific toxins, with 908 and 344 unique types, respectively (Fig. 1I). Correlation analysis between the expression of key annotated toxins and humidity levels revealed that the abundance of fibrinogenase, phospholipase, and phosphatase was significantly associated with humidity, while other tested toxins showed no significant correlation (Fig. 1J, Fig. S3).

### 3.2. Divergent functional profiles of scorpion venoms from different humidity regions

Building on the transcriptomic differences, we aimed to characterize the venom toxicity of these two scorpion species at the functional level. To enable a meaningful comparative toxicity assessment, we selected venom samples from regions with a comparable number of annotated toxins: AH and LN (representing *U. mordax*) and SX and NX (representing *M. martensii*). H9C2 cardiomyocytes were treated with venoms from these four regions for 6 h. Flow cytometry analysis revealed that venoms from the SX and NX induced a significantly higher apoptotic rate than those from LN and AH (Fig. 2A-C). It is noteworthy that there are inherent differences in venoms between the two distinct scorpion species. However, whether the geographical environmental conditions in which these different species inhabit also act as one of the factors driving such venom toxicity differences remains to be confirmed.

**Fig. 2.**
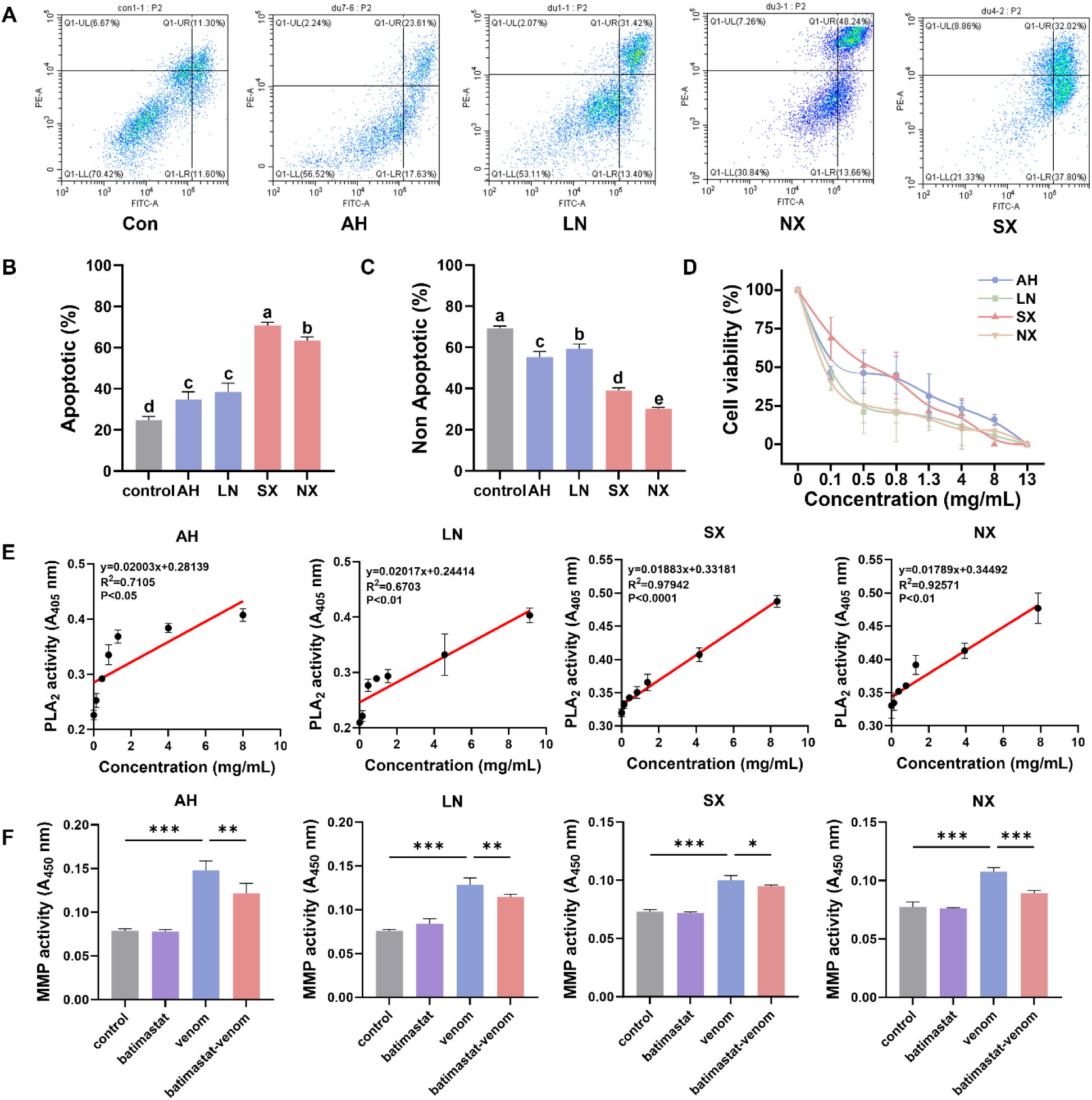
The effects of different environmental humidities on the toxicity and enzyme activity of scorpion venoms. **(A)** Flow cytometry plots depicting apoptosis of scorpion toxin cells in the AH, LN, NX, and SX regions (n=4, p<0.05); **(B-C)** Statistical plot of the effect of scorpion toxins on apoptosis and non-apoptosis in AH, LN, NX, and SX regions (n=4, p<0.05); **(D)** The impact of scorpion toxins in the AH, LN, NX, and SX regions on cell viability (n=3, p<0.05); **(E)** Compare the PLA_2_ activity of scorpion venom in AH, LN, SX and NX regions (n=3, p<0.05); **(F)** Compare the MMP activity of scorpion venom in AH, LN, SX and NX regions (n=3, p<0.05). Statistical significance among multiple groups was analyzed using one-way ANOVA, followed by Tukey’s HSD test for pairwise comparisons. Significant differences (p<0.05) are indicated using lowercase letters (a, b, c, etc.), groups that do not share a letter are significantly different.

Corroborating these findings, cell viability assays showed a concentration-dependent decrease in survival across all venoms, with complete cell death occurring at a concentration of 13 mg/mL (Fig. 2D). To gain mechanistic insight into the functional variation of venoms, we evaluated the activity of two key venom enzymes: phospholipase A_2_ (PLA_2_) and metalloprotease (MMP). PLA₂ activity increased consistently with venom concentration in all samples. Notably, venoms from the AH and LN exhibited significantly higher PLA_2_ activity than those from SX and NX (Fig. 2E). For the MMP activity assays, we included the broad-spectrum MMP inhibitor batimastat as a positive control to confirm that the detected activity was specifically attributable to scorpion venom-derived MMPs and not to cellular or non-specific proteases. Batimastat treatment markedly inhibited MMP activity across all venom samples, verifying the assay specificity. Similar to the PLA_2_ results, venoms from AH and LN also showed significantly higher MMP activity than those from SX and NX (Fig. 2F).

### 3.3. Identification and structural characterization of two novel calcins

A comprehensive toxin screening was performed across 8 regions, leading to the identification of two novel peptides. Phylogenetic tree analysis revealed that both peptides belong to the calcin family, and thus they were designated as M1 and M2. (Fig. 3A). M1 was specifically identified in *M. martensii* from the SX and XJ regions, while M2 was predominantly found in *U. mordax* from Tibet, AH, LN, and MM. The expression levels of both calcins were strongly associated with ambient humidity. Notably, M2 was enriched in all four high-humidity regions, though its expression level in Tibet (5.51 FPKM) was considerably lower than in other high-humidity areas. In contrast, M1 was only detected in two low-humidity regions and at significantly lower expression levels (Fig. 3B). Consistent with the calcin family, both M1 and M2 consist of 33 amino acids in their sequences and are typical basic hydrophilic peptides (Fig.3C). M1 contains four acidic and twelve basic amino acids, featuring five novel residues (Arg1, Lys4, Arg14, Asn19, Val28) not previously reported in calcins. M2 contains three acidic and ten basic amino acids, with Gly2 being a unique substitution (Fig. 3D).

**Fig. 3.**
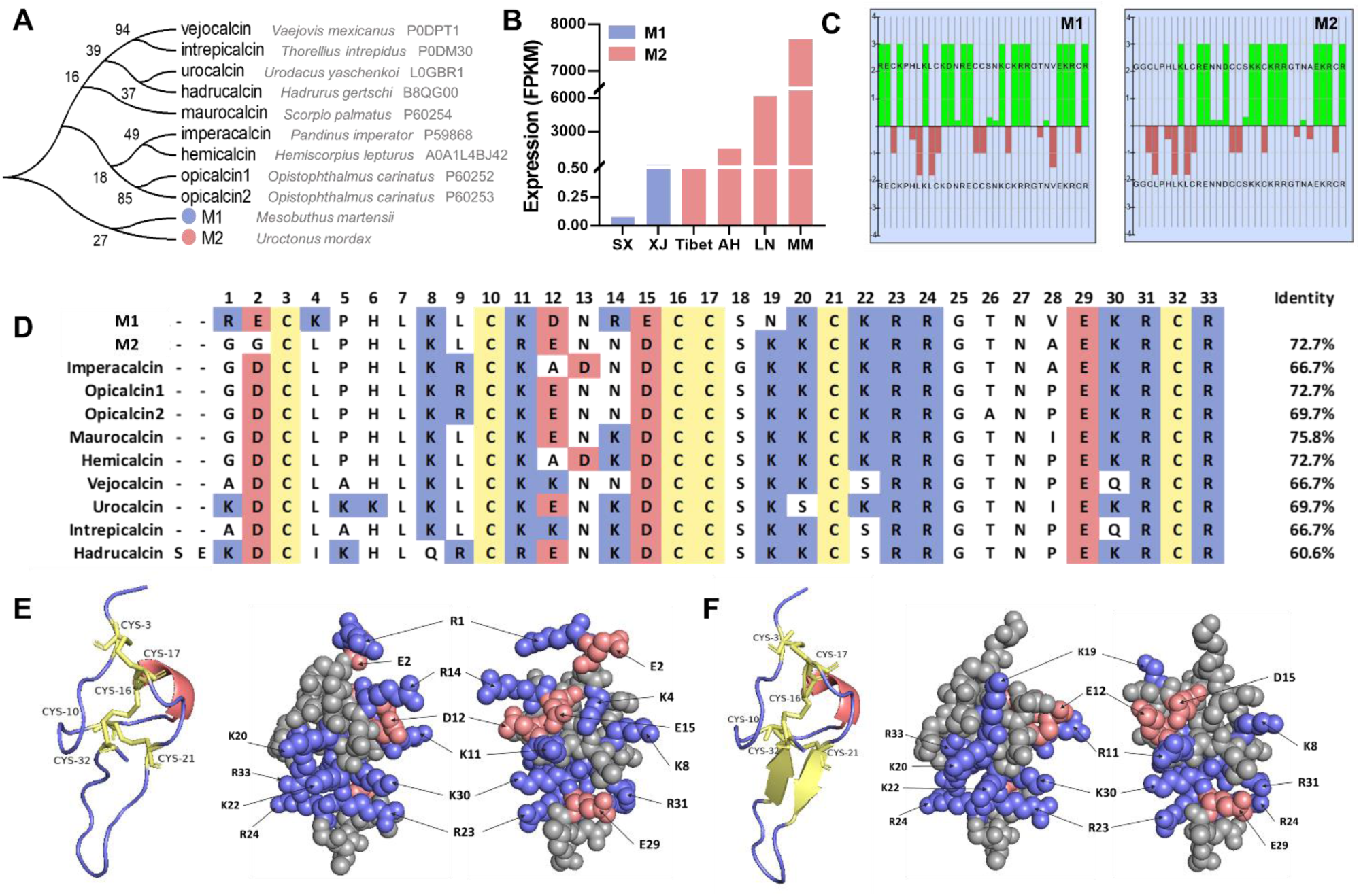
Discovery of two novel calcins and their structures. **(A)** Evolutionary analysis of M1 and M2 with known calcins; **(B)** Expression levels of M1 and M2 in scorpions from SX, Tibet, XJ, AH, LN and MM; **(C)** Hydrophobicity plot of M1 and M2. The ratio of hydrophilic residues (green) of M1 and M2 are between 61% and 52%, indicating high solubility in water for two novel calcins; **(D)** Sequence alignment among M1/M2 and other calcins. The positively charged residues lysine (K) and arginine (R), negatively charged residues aspartic acid (D) and glutamic acid (E), and the disulfide bond-forming cysteine (C) are highlighted by the colors blue, red, and yellow, respectively; **(E-F)** Spatial structure model of M1 and M2. α-helix, β-sheet, loop and disulfide bonds are marked in red, yellow and blue in the cartoon models. The basic amino acids, acidic amino acids and neutral amino acids are represented in blue, red and gray respectively in the sphere models.

Structurally, both M1 and M2 are predicted to adopt a globular peptide conformation, which is stabilized by a canonical inhibitor cystine knot (ICK) motif formed by the pairwise pairing of six cysteines in their sequences to generate three disulfide bonds. The first (Cys3-Cys17) and second (Cys10-Cys21) disulfide bonds create a loop, through which the third (Cys16-Cys32) disulfide bond passes, thus fashioning a knot. This structure confers stability and is typical of peptides that target ion channels. At the secondary structure level, M2 is characterized by an antiparallel β-sheet and one α-helix, while M1 contains only an α-helix (Fig. 3E and F). In both peptides, basic residues (e.g., 8KLCK11, 20KCKRR24, 30KRCR33 in M2) cluster on one molecular surface, creating a pronounced electrostatic polarity indicative of potential target interaction sites.

### 3.4. Differential binding and subconductance induction of RyR1 by two novel calcins

Molecular docking revealed that M1 could form eighteen pairs of interactions with RyR1, whereas M2 formed only eight. Except for Arg1, Lys11 and Lys20, all basic residues in the M1 sequence can form the hydrogen bonding interactions with RyR1. In M2, only five basic residues, Lys8, Lys20, Arg23, Arg31 and Arg33 can form the hydrogen bonding interactions with RyR1. Lys8, Lys20, Arg31 and Arg33 are conserved in both calcins and can form van der Waals force interaction with acidic residues on RyR1. However, compared to most members of the calcin family, only the fourth residue of M1 mutating from Leu to Lys. The remaining interacted residues, K22, R24 (M1), and R23 (M2) are also contained in the conserved cluster of calcin, forming hydrogen bonds (HBs) with the receptor (Fig. 4A-B, Table S1).

**Fig. 4.**
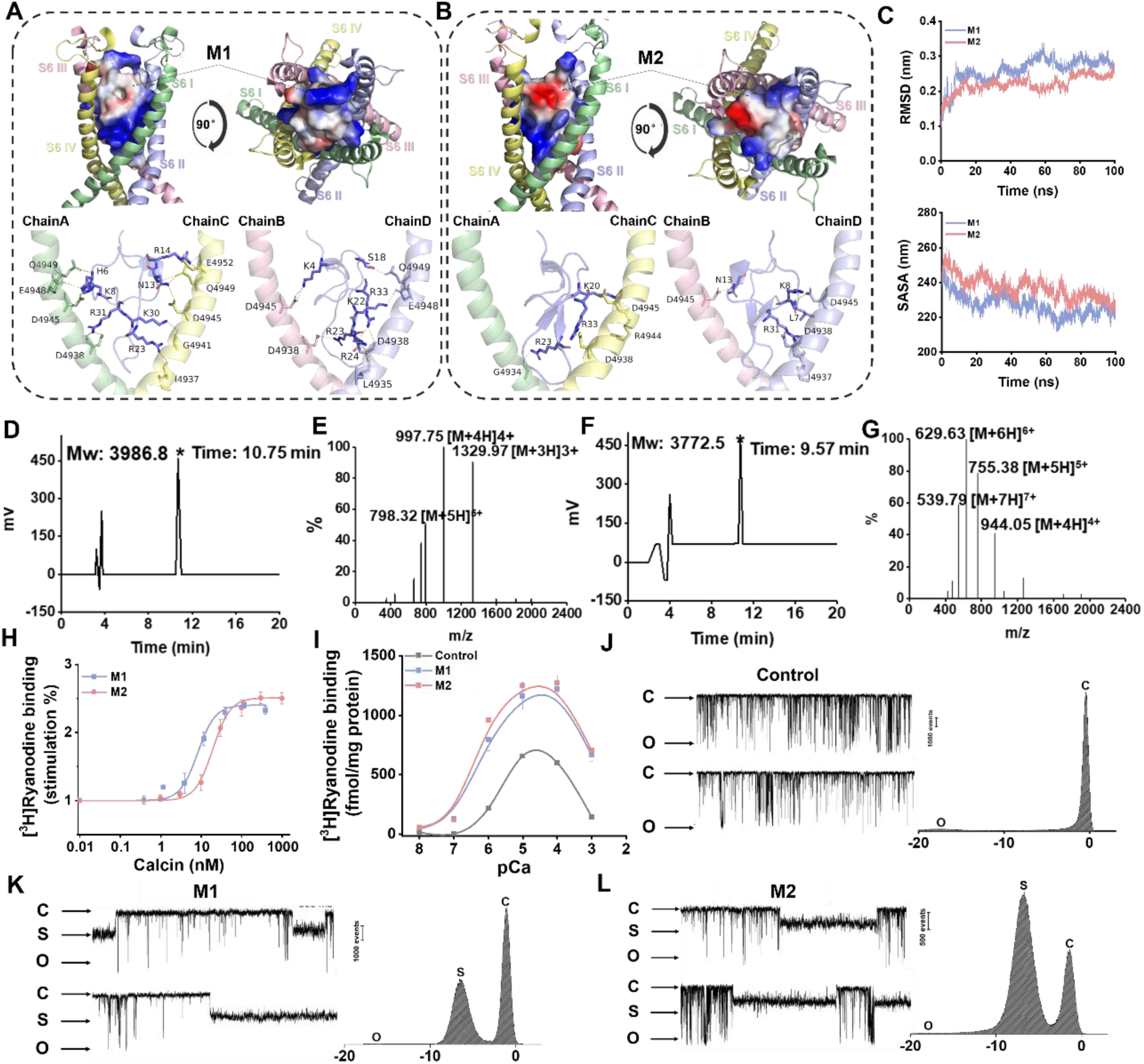
Synthesis, identification and activity detection of two novel calcins. **(A-B)** Details of residues in the interaction of M1 and M2 with S6 helices of RyR1 (PDB id: 7T65), all of which are bound by hydrogen bonds, and the bond length between the receptor and ligand is connected by yellow dashed line; **(C)** RMSD and SASA fluctuation curves of M1 and M2 docking with S6 helices in 100 ns molecular dynamics simulations; **(D-E)** Synthesis and Identification of M1; **(F-G)** Synthesis and Identification of M2; **(H)** Binding affinity of M1 and M2 (mean ± SEM, n=3); **(I)** Ca^2+^ sensitivity of [^3^H]ryanodine binding affected by M1 and M2 at 116 nM and 296 nM, respectively (mean ± SEM, n=3); **(J-L)** Single channel recording of control, M1 and M2: representative single RyR1 channel in the absence of calcin is shown as control; The M1 and M2 were separately introduced into the cis (cytosolic) compartment at the specified concentration.

The molecular dynamics simulation results revealed that the RMSD values of the M1 and M2 complex with RyR1 exhibited an initial increase from 0.1 to 0.2-0.3 nm within the first 20 ns, while remaining below the threshold of 0.4 nm, thereby indicating the acceptability of both complex conformation models. Concurrently, a gradual decrease in SASA value suggested a progressive compaction of the composite structure (Fig. 4C). The Gibbs free energy landscape (GFEL) and RMSF were employed to further investigate the conformational stability of calcins-RyR1 at its lowest energy state, as well as to compare the initial and final shifts in conformation. The results revealed that the binding of M1 and M2 to RyR1 exhibited an energy well, indicating that both novel calcins can form stable binding with RyR1 (Fig. S4A-C). In addition, dynamic simulations showed that the S6 helices exhibited a significant positional shift during the interaction of M1 and M2 with RyR1, as evidenced by comparing the first frame and the last frame of RMSF conformations extracted from the trajectory. This suggests that the interaction induced an asymmetric alteration in the height of the four S6 helices, thereby widening the pore gap (Fig. S4D-E).

The target peaks of M1 (10.75 min) and M2 (9.57 min) were collected from HPLC analysis, followed by identification using mass spectrometry. The molecular weights of the products were determined to be 3986.8 Da and 3772.5 Da, respectively (Fig. 4D-G). Finally, we obtained M1 and M2 with a purity of ≥ 95%. Both M1 and M2 were found to stimulate [^3^H]ryanodine binding of RyR1 with different affinities (apparent Kd: 8.2 nM (M1) vs. 19.3 nM (M2)) but similar potency (maximal stimulation normalized to control, ≈2.5) (Fig. 4H). Consistent with the results from other calcins, both of the two novel calcins increased [^3^H]ryanodine binding to RyR1 at low Ca^2+^ concentrations, while decreasing the maximum amount of [^3^H]ryanodine binding at higher Ca^2+^ concentrations. The maximal values of the binding for these two calcins were at pCa5 to 4 with similar amplitude (Fig. 4I). In order to further identify the functional properties of calcins, RyR1 channels were incorporated into planar lipid bilayers for the examination of calcins on single-channel gating. Consistent with prior findings, the application of M1 and M2 (each at 300 nM) led to a distinct response in RyR1, characterized by the emergence of a reversible, long-lasting, transient subconductance state. However, the sub-states induced by M1 and M2 corresponded to 0.33 and 0.26 of the full-conductance levels, respectively, which were close to those previously reported for Urocalcin (0.55) and Hadrucalcin (0.35)(Xiao et al., 2016) (Fig. 4J-L). In all, the two calcins can strongly stimulate the [^3^H]ryanodine binding to RyR1 with high affinities and induce the long-lasting subconductances that were similar to those of other known calcins.

### 3.5. Lys4 residue plays a pivotal role in determining the disparity in the activity of the two novel calcins

To elucidate the critical amino acid residues responsible for the differential binding affinity of M1 and M2 towards RyR1, we conducted a comprehensive mutagenesis analysis. We first performed computational alanine scanning and mutual mutation simulations at equivalent positions in M1 and M2 (Fig. 5A). The change in binding free energy (ΔiG) predicted that E2G and K4L mutations in M1 would significantly destabilize the binding, whereas mutations K11R, E15D, and N19K were predicted to stabilize it. In M2, most mutations at the equivalent sites exhibited opposite effects on ΔiG compared to M1, except for V28A (Fig. 5B). Based on these predictions and considering the pivotal role of electrostatic interactions, we synthesized two M1 mutants: E2G and K4L (Fig. 5C). Radioligand binding assays revealed that the K4L mutation significantly weakened the binding affinity, increasing the apparent Kd value to 22.0 nM, which is nearly identical to that of wild-type M2 (19.3 nM) (Fig. 5D). In contrast, the E2G mutation did not significantly alter the affinity for RyR1. This indicates that Lys4 is a primary determinant underlying the higher binding affinity of M1 compared to M2.

**Fig. 5.**
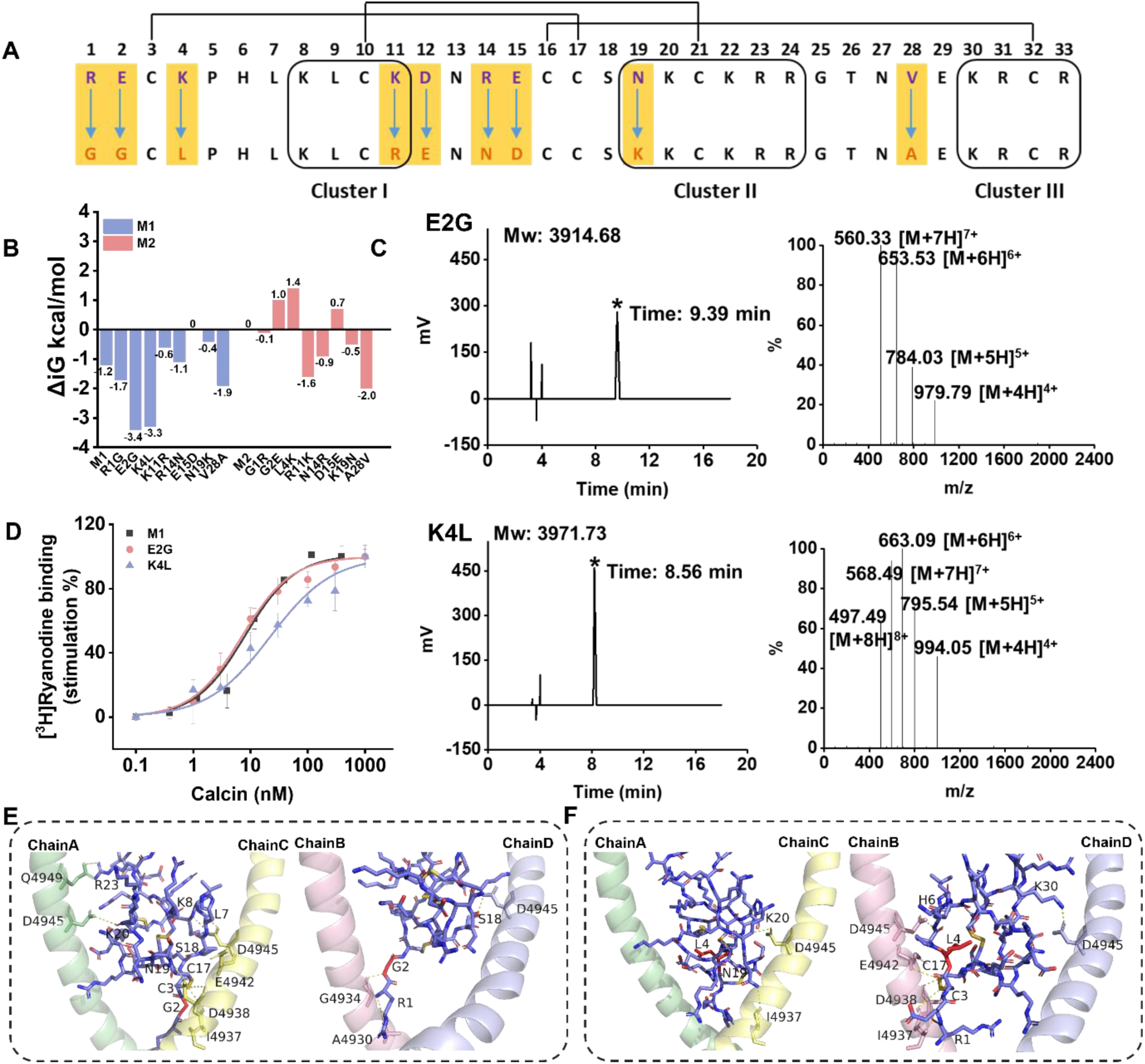
Comparison of binding affinity and activity between mutant peptide analogs of M1 and RyR1. **(A)** Mutational substitution of different residues at the same position in M1 and M2; **(B)** Comparison of free energy values of M1 and M2 mutant peptides (blue: M1 free energy value; red: M2 free energy value); **(C)** Synthesis and Identification of mutant E2G and KEL; **(D)** Stimulation of [^3^H] ryanodine binding to RyR1 by M1 mutants including E2G and K4L (n=3); **(E-F)** Details of residues in the interaction of E2G and K4L with RyR1, all of which are bound by hydrogen bonds, and the bond length between the receptor and ligand is connected by yellow dashed lines, mutated amino acids are represented by red sticks.

To understand the structural basis of this finding, we analyzed the binding modes of the mutants. The E2G mutant formed twelve interaction pairs with RyR1. The removal of the negatively charged Glu2 side chain reduced steric hindrance and electrostatic repulsion, allowing Cys3 to form multiple hydrogen bonds with the S6 helices (Fig. 5E, Table S2). However, this did not translate into higher affinity, suggesting that the beneficial effect of reduced repulsion was offset by the loss of other interactions. In contrast, the K4L mutation not only eliminated a key favorable electrostatic interaction but also drastically reduced the number of hydrogen bonds between the peptide and RyR1 (Fig. 5F, Table S2). The long side chain of Lys4 in wild-type M1 penetrates into the central channel domain of RyR1, which is rich in acidic residues and forms strong charge-charge interactions. Substituting Lys with the neutral Leu abolishes this interaction and reduces the overall basicity of the peptide, thereby explaining the significant loss of affinity. In summary, our integrated computational and experimental approach identifies Lys4 as a critical residue responsible for the enhanced binding affinity of M1 over M2. This difference primarily stems from the additional electrostatic and hydrogen-bonding interactions mediated by this residue.

## 4. Discussion

Scorpion envenomation constitutes a significant global health burden, yet the same venoms represent a rich source of bioactive peptides with considerable therapeutic potential(Bahloul, Regaieg et al., 2017, Dabas, 2019, Reis, Zoccal et al., 2019). While interspecific variation in venom composition is well established, the extrinsic factors driving intraspecific variability and functional adaptation remain poorly characterized(Nystrom, Ward et al., 2019). Through integrated geographical sampling, transcriptomic analysis, functional assays, and structural modeling, this study identified two novel calcins from two distinct scorpion species, namely *Uroctonus mordax* and *Mesobuthus martensii*, and clarified the molecular basis underlying their functional differences. Furthermore, this study proposes that although there are inherent differences in venom toxicity among different scorpion species, geographical environmental conditions may still serve as external factors driving variations in venom toxicity and enzymatic activity.

Our transcriptomic analyses revealed biogeographical variation in venom gland expression profiles between and within species. Notably, we observed that venom cytotoxicity was higher in samples from SX and NX (*M. martensii* from low-humidity regions), whereas enzymatic activities (PLA_2_ and MMP) were elevated in venoms from AH and LN (*U. mordax* from high-humidity regions). Regarding this functional variation, where scorpion venoms exhibit high cytotoxicity alongside low activity of key enzymes, it may stem from two interacting factors. Primarily, this variation is ascribed to inherent interspecific differences dictated by the phylogenetic background of the scorpion species(Anandhan Sujatha, Gopalakrishnan et al., 2024, Hudefe, Álvarez et al., 2024). In addition, geographical variation shaped by environmental selection pressure may also serve as a contributing factor(Mabunda, Zinyemba et al., 2024, Rodríguez-Ravelo, Coronas et al., 2013, Yu, Yu et al., 2020). This suggests a climate-driven functional trade-off: scorpions may evolve venoms to have heightened cytolytic activity in low-humidity environments and potentially to maximize prey immobilization efficiency under resource-limited conditions, while favoring enzymatic components in high-humidity environments, possibly for enhanced digestion or antimicrobial defense(Díaz, Rivera et al., 2019, Goudet, Chi et al., 2002, Salazar, Samudio et al., 2025, Samudio, Hernández-Ortiz et al., 2025). This pattern of environmental adaptation is not unique to scorpions. For instance, environmental factors can also exert regulatory effects on snake venom toxicity, pointing to a parallel adaptive strategy in which such factors fine-tune venom composition to maximize its ecological functionality(Chippaux, Saz-Parkinson et al., 2013, Keyler, Gawarammana et al., 2013). These findings align with growing evidence that abiotic factors can profoundly shape venom composition, possibly through selective pressures acting on biosynthetic pathways or gene regulation mechanisms(Barkan, Chevalier et al., 2019, Soltan-Alinejad, Alipour et al., 2022).

Beyond overall venom activity, we identified two novel calcins whose distribution and expression patterns revealed an interplay between species identity and environmental adaptation. Furthermore, within each species, calcin expression levels exhibited a significant association with habitat humidity. Specifically, M2 expression was highest in *U. mordax* populations from high-humidity regions (though constrained in the high-altitude environment of Tibet), while M1 was expressed at low levels in *M. martensii* populations from arid, low-humidity regions. Both peptides exhibit the conserved structural of calcins, including an ICK motif, a high proportion of basic residues, and a hydrophilic molecular surface(Hua, Yao et al., 2024). These features are critical for RyR targeting and channel modulation. Functionally, consistent with other calcin family members, both M1 and M2 activated RyR1 and induced a subconductance state. However, M1 exhibited significantly higher binding affinity (Kd=8.2 nM vs. 19.3 nM), highlighting functional divergence within the family. Through integrated computational and experimental mutagenesis, we identified Lys4 in M1 as a pivotal residue responsible for this enhanced affinity. Substitution to Leu (K4L) reduced M1’s binding affinity to a level comparable to wild-type M2, underscoring the importance of electrostatic interactions with acidic residues within the RyR1 central channel. This residue-specific insight provides a mechanistic basis for the differential activity of these peptides and illustrates how subtle sequence variations can underlie ecological adaptations(Haji-Ghassemi, Chen et al., 2023, Yao, Hua et al., 2024). The geographical structuring of toxin expression, particularly the positive correlation between humidity and certain toxin groups, which suggests that environmental factors may act as selective agents fine-tuning venom composition(Koludarov, Velasque et al., 2023, von Reumont, Dutertre et al., 2022). This adaptive tuning may optimize venom efficacy under specific climatic conditions, although the exact ecological drivers (e.g., prey spectrum, physiological constraints) warrant further investigation(Valdez-Cruz, Dávila et al., 2004, Webber, Gibbs et al., 2015).

Scorpions have long been used in traditional Chinese medicine, and our identification of two novel RyR-targeting calcins with differential activity underscores the potential of environmentally informed venom screening for discovering ion channel modulators with refined therapeutic properties(Haji-Ghassemi et al., 2023, Xiao et al., 2016). Although the eight sampled regions encompass notable climatic diversity, the relatively small number of geographical samples and the focus on only two species limit the generalizability of our conclusions. In particular, the confounding effects of species identity and regional climate cannot be fully disentangled here. *Uroctonus mordax* and *Mesobuthus martensii* may possess inherent differences in venom composition independent of environmental influence(Santibáñez-López et al., 2019, Santibáñez-López, Kriebel et al., 2018). Nevertheless, despite these phylogenetic and sampling constraints, we observed a consistent and strong correlative relationship between environmental humidity and multiple layers of venom variation, including the number of assembled unigenes, the expression of specific toxins (such as fibrinogenase, phospholipase, and phosphatase), and the geographical distribution of the two novel calcins (M1 and M2). This recurring pattern suggests that humidity likely acts as an important environmental modulator of venom composition, possibly superimposing its effects on underlying phylogenetic determinants(Cid-Uribe, Santibáñez-López et al., 2018, Di, Qiao et al., 2022). We further identify Lys4 in M1 as a critical residue governing calcin-RyR1 binding affinity, unveiling a molecular mechanism for functional divergence between these novel peptides. These findings enhance our understanding of ecological adaptation in venomous organisms and highlight the potential of environment-guided biodiscovery for identifying functionally refined bioactive peptides with therapeutic promise.

## 5. Conclusions

This study demonstrates that environmental humidity is a key driver shaping the molecular diversity of scorpion venoms across transcriptional, functional, and structural levels. Through a comprehensive analysis of specimens from diverse climatic zones in East Asia, we revealed that venom gland transcriptomes particularly the number of assembled unigenes, which positively correlated with regional humidity. Furthermore, we found that the expression of several key toxins, including fibrinogenase, phospholipase, phosphatase, and two novel calcin peptides, is influenced by humidity conditions. Functionally, both novel calcins (M1 and M2) target RyRs, inducing a characteristic subconductance state, yet they exhibit significant differences in binding affinity. We identified Lys^4^ as the critical residue responsible for this functional divergence, providing a mechanistic understanding of how subtle sequence variations can underlie ecological adaptations. Our findings not only illuminate the intricate evolutionary strategies employed by scorpion venoms to adapt to their environments but also highlight the potential of environment-guided bioprospecting for the discovery of functionally refined bioactive molecules with therapeutic potential.

## CRediT authorship contribution statement

**Xiaoyu Hua:** Conceptualization, Investigation, Writing—review & editing, Writing—original draft; **Zhixiao Yang:** Formal Analysis, Writing—review & editing; **Li Xiao:** Methodology, Software, Writing—review & editing; **Songyu Gao:** Visualization, Methodology; **Fengling Yang:** Resources, Visualization; **Yi Wang:** Visualization, Validation; **Jianguo Wang:** Investigation; **Carmen R. Valdivia:** Supervision; **Wei Chen:** Methodology, Visualization; **Wengeng Jiang, He Sun, Diguyan Wu:** Validation, Visualization; **Marina Pozzolini:** Methodology, Visualization; **Lingxin Chen:** Supervision, Writing—review & editing; **Héctor H Valdiva:** Conceptualization, Resources, Supervision; **Liang Xiao:** Conceptualization, Funding Acquisition, Supervision, Writing—review & editing.

## Acknowledgement

This work was funded by 2023 National Key R&D Program of China (the Ministry of Science and Technology Sino US International Cooperation Project) [Grant. no: 2023YFE0117800], and University-level basic medical research Project of Naval Medical University [Grant. no:2024QN008].

